# Broad and strong memory CD4^+^ and CD8^+^ T cells induced by SARS-CoV-2 in UK convalescent COVID-19 patients

**DOI:** 10.1101/2020.06.05.134551

**Authors:** Yanchun Peng, Alexander J. Mentzer, Guihai Liu, Xuan Yao, Zixi Yin, Danning Dong, Wanwisa Dejnirattisai, Timothy Rostron, Piyada Supasa, Chang Liu, Cesar Lopez-Camacho, Jose Slon-campos, Yuguang Zhao, Dave Stuart, Guido Paeson, Jonathan Grimes, Fred Antson, Oliver W. Bayfield, Dorothy EDP. Hawkins, De-Sheng Ker, Lance Turtle, Krishanthi Subramaniam, Paul Thomson, Ping Zhang, Christina Dold, Jeremy Ratcliff, Peter Simmonds, Thushan de Silva, Paul Sopp, Dannielle Wellington, Ushani Rajapaksa, Yi-Ling Chen, Mariolina Salio, Giorgio Napolitani, Wayne Paes, Persephone Borrow, Benedikt Kessler, Jeremy W. Fry, Nikolai F. Schwabe, Malcolm G Semple, Kenneth J. Baillie, Shona Moore, Peter JM Openshaw, Azim Ansari, Susanna Dunachie, Ellie Barnes, John Frater, Georgina Kerr, Philip Goulder, Teresa Lockett, Robert Levin, Oxford Immunology Network Covid-19 Response T cell Consortium, Richard J. Cornall, Chris Conlon, Paul Klenerman, Andrew McMichael, Gavin Screaton, Juthathip Mongkolsapaya, Julian C. Knight, Graham Ogg, Tao Dong

## Abstract

COVID-19 is an ongoing global crisis in which the development of effective vaccines and therapeutics will depend critically on understanding the natural immunity to the virus, including the role of SARS-CoV-2-specific T cells. We have conducted a study of 42 patients following recovery from COVID-19, including 28 mild and 14 severe cases, comparing their T cell responses to those of 16 control donors. We assessed the immune memory of T cell responses using IFNγ based assays with overlapping peptides spanning SARS-CoV-2 apart from ORF1. We found the breadth, magnitude and frequency of memory T cell responses from COVID-19 were significantly higher in severe compared to mild COVID-19 cases, and this effect was most marked in response to spike, membrane, and ORF3a proteins. Total and spike-specific T cell responses correlated with the anti-Spike, anti-Receptor Binding Domain (RBD) as well as anti-Nucleoprotein (NP) endpoint antibody titre (p<0.001, <0.001 and =0.002). We identified 39 separate peptides containing CD4^+^ and/or CD8^+^ epitopes, which strikingly included six immunodominant epitope clusters targeted by T cells in many donors, including 3 clusters in spike (recognised by 29%, 24%, 18% donors), two in the membrane protein (M, 32%, 47%) and one in the nucleoprotein (Np, 35%). CD8+ responses were further defined for their HLA restriction, including B*4001-restricted T cells showing central memory and effector memory phenotype. In mild cases, higher frequencies of multi-cytokine producing M- and NP-specific CD8^+^ T cells than spike-specific CD8^+^ T cells were observed. They furthermore showed a higher ratio of SARS-CoV-2-specific CD8^+^ to CD4^+^ T cell responses. Immunodominant epitope clusters and peptides containing T cell epitopes identified in this study will provide critical tools to study the role of virus-specific T cells in control and resolution of SARS-CoV-2 infections. The identification of T cell specificity and functionality associated with milder disease, highlights the potential importance of including non-spike proteins within future COVID-19 vaccine design.

## Introduction

COVID-19 is caused by a recently emerged RNA beta coronavirus, Severe Acute Respiratory Syndrome coronavirus-2 (SARS-CoV-2). Whilst the majority of COVID-19 infections are relatively mild, with recovery typically within two to three weeks ^1, 2^, a significant number of patients develop severe, often fatal illness, which is postulated to be related to both an overactive immune response and viral induced lung pathology ^3, 4^. The role of adaptive T cell immune responses in disease pathogenesis and longer-term protective immunity is currently poorly defined but essential to inform potential therapeutic interventions and enhance vaccine design.

Currently, there are several ongoing vaccine trials, but uncertainties abound and it is unknown whether they will provide long lasting protective immunity. Most vaccines are aimed to induce humoral immunity to the SARS-CoV-2 spike protein, but it is not yet known if this will be sufficient to induce full protective immunity to SARS-CoV-2. A recent study assessed the immune response induced by an adenovirus-vectored vaccine ChAdOx1 nCoV-19, encoding the spike protein of SARS-CoV-2 and showed protection from SARS-CoV-2-associated pneumonia in rhesus macaques, but without inducing sterilising immunity ^5^. A similar observation was also made by Yu *et al* when using a DNA vaccine candidate expressing different forms of the SARS-CoV-2 spike (S) protein in a rhesus macaque model^6^. Another study from China in humans assessed safety, tolerability, and immunogenicity of a recombinant adenovirus type-5 (Ad5) vectored COVID-19 vaccine, which induced both T cell responses and neutralizing antibody responses to SARS-CoV-2 specifically^7^. Studying natural immunity to the virus, especially the role of SARS-CoV-2-specific T cells is critical to fill the current knowledge gaps for improved vaccine design.

For primary virus infection, it typically takes 7-10 days for adaptive T cell immune responses to be primed and expanded in order to control the virus^8^. This coincides with the typical period of time in which COVID-19 patients to either recover or develop severe illness. There is a median incubation time of 4-7 days in patients before patients developing symptoms and a further 7-10 days before a minority cases progress to severe disease requiring intensive care unit admission and/or mechanical ventilation occurs in a minority of cases ^9^. Such a pattern of progression raises the possibility that relative immunosuppression contributes to SARS-CoV-2 viral persistence and COVID-19 mortality, whereas adaptive immune responses are usually protective in the majority of the population.

While T cells are not thought to prevent the establishment of infection, there is good evidence in mice and humans that T cells provide partial protection against many acute viral infections, such as influenza, by promoting viral clearance and reducing the severity of symptoms ^1, 10^. Single cell analysis of bronchoalveolar lavage fluid (BALF) from COVID-19 patients revealed clonal expansions of CD8^+^ T cells in mild but not severe patients, suggesting that the presence of adaptive T cell responses may be protective in SARS-CoV-2 infection ^11^.

Evidence supporting a role for T cells in COVID-19 pathogenesis is currently incomplete and conflicting. Some reports indicate that an overreaction of T cells with high cytotoxicity that causes lung injury^3^ rather than a lack of a T cell response; one post-mortem study suggested that direct infection of ACE2-expressing macrophages could lead to activation-induced T cell death (AICD)^12^. In contrast a lack of functional T cells may also contribute to severe disease since lymphocytopenia in COVID-19 patients, especially in older and critically ill cases, may be the cause of functional exhaustion^13^.

To date there have been few studies analysing SARS-CoV-2-specific T cell responses and their role in disease progression ^14^. Using HLA-class I and II predicted “Mega” peptide pools, Grifoni *et al* evaluated CD4^+^ and CD8^+^ T cell responses in non-hospitalised convalescent subjects. They found all recovered subjects established CD4^+^ responses and 70% established CD8^+^ memory responses to SARS-CoV-2^15^. SARS-CoV-2-specific CD4^+^ T cell responses were also frequently observed in unexposed subjects in their study, suggesting the possibility of pre-existing cross-reactive immune memory to seasonal coronaviruses.

In the present study, the overall and immunodominant SARS-CoV-2–specific memory T cell response in subjects who had recovered from COVID-19 were evaluated *ex vivo* using overlapping peptides spanning the full proteome of the SARS-CoV-2, excepting only ORF-1. Potential epitopes were identified using two-dimensional matrix peptide pools and further evaluated for CD4^+^ and CD8^+^ subset involvement. The function of SARS-CoV-2-specific T cells specific for dominant T cell responses were evaluated by intracellular cytokine production profiles, and the HLA restriction of the dominant CD8^+^ T cell responses was defined using *ex vivo* and *in vitro* cultured short-term T cell lines. Broad, and frequently strong, SARS-CoV-2 specific CD4^+^ and CD8^+^ T cell responses were seen in the majority of convalescent patients, with a greater proportion of CD8+ T cell responses in mild cases, together with higher frequencies of multi-cytokine producing M- and NP-specific CD8+ T cells.

## Materials and methods

Patients were recruited from the John Radcliffe Hospital in Oxford, UK, between March and May 2020 by identification of patients hospitalised during the SARS-COV-2 pandemic and recruited into the Sepsis Immunomics and ISARIC Clinical Characterisation Protocols. Patients were sampled at least 28 days from the start of their symptoms. Unexposed healthy adult donor samples were used from unrelated studies undertaken between 2017-early 2019, and from sero-negative individuals in 2020. Written informed consent was obtained from all patients.

### Clinical definitions

All patients were confirmed to have a test positive for SARS-CoV-2 using reverse transcriptase polymerase chain reaction (RT-PCR) from an upper respiratory tract (nose/throat) swab tested in accredited laboratories. The degree of severity was identified as mild, severe or critical infection according to recommendations from the World Health Organisation. Severe infection was defined as COVID-19 confirmed patients with one of the following conditions: respiratory distress with RR>30/min; blood oxygen saturation<93%; arterial oxygen partial pressure (PaO2) / fraction of inspired O2 (FiO2) <300mmHg; and critical infection was defined as respiratory failure requiring mechanical ventilation or shock; or other organ failures requiring admission to ICU. Since the Severe classification could potentially include individuals spanning a wide spectrum of disease severity spanning patients receiving nasal cannula through to non-invasive ventilation we also calculated the SaO2/FiO2 ratio at the height of patient illness as a quantitative marker of lung damage. This was calculated by dividing the oxygen saturation (as determined using a bedside pulse oximeter) by the fraction of inspired oxygen (21% for ambient air, 24% for nasal cannulae, 28% for simple face masks and 28, 35, 40 or 60% for Venturi face masks or precise measurements for non-invasive or invasive ventilation settings).

### Synthetic peptides

A total of 423 15- to 18-mer peptides overlapping by 10 amino acid residues and spanning the full proteome of the SARS-CoV-2 except ORF-1 (253 spike, 29 M, 9 E, 35 ORF3a, 7ORF6, 15 ORF7a, 16 ORF8, 59 NP) were designed using the Los Alamos National Library web-based software PeptGen (http://www.hiv.lanl.gov/content/sequence/PEPTGEN/peptgen.html) and synthesized (purity >75%; Proimmune).

A set of previously defined 27 SARS epitopes ^16^ were also synthesised (Table 2), and 19 tetramer and pentamers with those peptides in complex with appropriate Class I and II HLAs were provided by Proimmune.

Pools of Cytomegalovirus (CMV),Epstein-Barr cirus (EBV) and influenza virus specific epitope peptides and The human immunodeficiency viruses (HIV) gag were also used as positive and negative controls.

### 2-dimensional peptide matrix system

The overlapping peptides spanning the SARS-CoV-2 were assigned into a 2-dimensional matrix system in which each peptide was represented in 2 different peptide pools. Each peptide pool contains no more than 16 individual peptides. The first dimension of the peptide matrix system was designed so that peptides from different source proteins were separated into different pools. (Supplemental Table 1).

### *Ex vivo* ELISpot assay

IFN-γ ELISpot assays were performed using either freshly isolated or cryopreserved PBMCs as described previously. No significant difference was observed between responses generated by fresh or cryopreserved PBMCs as described previously^17, 18^.

Overlapping peptides were pooled such that the final concentration of each peptide used was 2 μg/mL and then added to 200,000 PBMCs per test for 16–18 h. To quantify antigen-specific responses, mean spots of the control wells were subtracted from the positive wells, and the results expressed as spot forming units (SFU)/10^6^ PBMCs. Responses were considered positive if results were at least three times the mean of the negative control wells and >25 SFU/10^6^PBMCs. If negative control wells had >30 SFU/10^6^ PBMCs or positive control wells (PHA stimulation) were negative, the results were excluded from further analysis.

### Determination of plasma binding to trimeric spike, RBD and NP by ELISA

MAXISORP immunoplates (442404; NUNC) were coated with 0.125 ug of StrepMAB-Classic (2-1507-001;iba), blocked with 2% skimmed milk in PBS for one hour and then incubated with 50 ul of 5 ug/ml soluble trimeric Spike 2 ug/ml or 2% skim milk in PBS. After one hour, 50 μL of serial two-fold dilutions of plasma, from 1:50 to 1:51200 in PBS containing 2% skimmed milk were added followed by ALP-conjugated anti-human IgG (A9544; Sigma) at 1:10,000 dilution. The reaction was developed by the addition of PNPP substrate and stopped with NaOH. The absorbance was measured at 405nm. Endpoint titers (EPTs) were defined as reciprocal plasma dilutions that corresponded to two times the average OD values obtained with mock. To determine EPTs to RBD and NP, immunoplates were coated with 0.125ug of Tetra-His antibody (34670; QIAGEN) followed by 2 ug/ml and 5 ug/ml of soluble RBD and NP, respectively.

### Intracellular cytokine staining (ICS) based T-Cell Assay

Confirmed epitope-containing peptides and positive peptide pools were characterized further for their CD4^+^ and CD8^+^ subset involvement and cytokine production pattern by intracellular cytokine staining as described previously^19, 20^. Briefly, cryopreserved PBMCs were thawed and rested overnight in R10 at 37°C. On the second day, the PBMCs were stimulated with pooled or individual peptides at a final concentration of 10 μg/mL per individual peptide for 1 h in the presence of 2 μg/mL monoclonal antibodies against human CD28 (BD Pharmingen) and CD49d (BD Pharmingen) then for an additional 5h with GolgiPlug (brefeldin A, BD), GolgiStop (monensin, BD) and surface stained with PE-anti-CD107a (BD Biosciences). For the FACS staining, dead cells were first labelled using LIVE/DEAD™ Fixable Aqua Dead Cell Stain Kit (Invitrogen). Finally, surface markers including BUV395-anti-CD3 (BD Biosciences), BUV737-anti-CD4 (BD Biosciences), PerCP-Cy5.5-anti-CD8 (BD Biosciences), BV510-anti-CD14 (Biolegend), BV510-anti-CD16 (Biolegend) and BV510-anti-CD19 (Biolegend) were stained. Cells were then washed, fixed with Cytofix/Cytoperm™ (BD Biosciences) and stained with PE-Cy7-anti-IFNγ (BD Biosciences), APC-anti-TNFα (eBioscience), BV421-anti-I-L2 (Biolegend). Negative controls with no peptide stimulation were run in parallel for each sample. Cytokine responses were background subtracted. All samples were acquired on BD LSR Fortessa (BD Biosciences) flow cytometer and analyzed using FlowJo™ v.10 software for Mac (Becton, Dickinson and Company; 2019). To determine the frequency of different response patterns based on all possible combinations, Boolean gate were created using IFN-γ, TNF-α and IL-2. The frequency of responding cells in the negative control (anti-CD28/anti-CD49d) were subtracted from that in the stimulated samples for each response individually prior to further analysis. Those which yielding negative values were set to 0.

### Pentamer phenotyping

Cryopreserved PBMCs were thawed as described above. A total of 1 × 10^6^ live PBMCs were labeled with peptide-MHC class I pentamer-PE (Proimmune, UK) and incubated for 15 min at 37°C. Dead cells were first labelled for FACS analysis using LIVE/DEAD™ Fixable Aqua Dead Cell Stain Kit (Invitrogen) and with CD3-BUV395, CD8-PerCP.Cy5.5 as well as a panel of antibodies for dumping, cell activation, differentiation and inhibitory markers : CD14-BV510 (Biolegend UK), CD16-BV510 (Biolegend UK), CD19-BV510 (Biolegend UK), CD28-BV711, CD27-APC-R700, HLA-DR-BB515, CD38-BUV737, CD45RA-APC-H7, PD-1-BV650, CD57-BV785 (Biolegend, UK) and NKG2A (R&D). Cells were then permeabilized with Perm/fix (BD, Oxford, UK) for 15 min and washed twice with 1× perm/washing buffer (BD) before being stained with Ki-67-BV421 and GranzymeB-PE-Cy7 (Biolegend, UK). Cells were subsequently washed twice with 1× perm/washing buffer and fixed in BD cellfix (BD Bioscience). All antibodies were from Becton Dickinson (BD, Oxford, UK) unless otherwise stated. Cell events were acquired on Fortessa X20 (BD Bioscience) and data files were analyzed using FlowJo software. Data were analyzed using a forward side scatter single cell gate followed by CD8 gating, then tetramer gating within the CD8^+^ population. These cells were then analyzed for percentage expression of a particular marker using unstained and overall CD8^+^ populations to determine where to place the gates. Single-color samples were run for compensation, and fluorescence minus one (FMO) control samples were also applied to determine positive and negative populations, as well as channel spillover.

### Generating short-term T cell lines

Short-term SARS-CoV-2-specific T cell lines were established as previously described ^18^. Briefly, 3 × 10^6^ to 5 × 10^6^ PBMCs were pulsed as a pellet for 1 h at 37°C with 10 μM of peptides containing T cell epitope regions and cultured in R10 at 2 × 10^6^ cells per well in a 24-well Costar plate. IL-2 was added to a final concentration of 100 U/ml on day 3.

### Statistical analysis

Statistical analysis was performed with IBM SPSS Statistics 25 and figures were made with GraphPad Prism 8. Chi-square tests were used to compare ratio difference between two groups. After testing for normality using Kolmogorov-Smirnov test, Independent-samples *t* test or Mann-Whitney U test was employed to compare variables between two groups. Correlations were performed via Spearman’s rank correlation coefficient. Statistical significance was set at P<0.05 and all tests were 2-tailed.

## Results

### Study subjects

42 individuals were recruited following recovery from COVID-19, including 28 mild cases and 14 severe cases. In addition, 19 control individuals sampled in the pre-COVID-19 season were studied in parallel. Supplementary Figure 1 shows the participant characteristics. No significant differences in gender or age were noted between mild and severe groups. The SaO2/FiO2 ratio in severe cases ranged from 4.3 (where 4.5 would be the estimate for an individual with mild disease breathing ambient air) to 1.6 with the patients with critical disease having an estimate of 0.8 (median in severe group 3.8).

### *Ex vivo* assessment of memory T cell responses specific to SARS-CoV-2

PBMCs were tested for responses to a panel of 423 overlapping peptides spanning all SARS-CoV-2 proteomes except ORF1 using *ex vivo* IFN-γ ELISpot assays. All overlapping peptides were placed into two 2-dimensional peptide matrix systems as described in Supplementary Table 1.

The majority of the participants exhibited SARS-CoV-2 memory T cell responses to at least one of the peptides. The overall distribution, magnitude and breadth of the IFN-γ responses against all SARS-CoV-2 virus protein peptides are shown in Figure 1. There is no correlation between the T cell responses and days post symptom development when sampled (Supplementary Figure 2). We did not detect any IFNγ-producing SARS-CoV-2-specific T cell responses in unexposed healthy volunteers, but were able to detect EBV, CMV and influenza virus specific T cell responses using pools of known T cell epitopes as controls. The breadth and magnitude of the T cell responses varied considerably between individuals. T cell responses were detected against epitopes distributed across a wide variety of virus proteins. A significantly higher magnitude (p=0.002) and broader breadth (p=0.010) overall T cell responses were observed in severe cases in comparison with mild cases, in particular responses to spike (magnitude/breadth, p=0.021/0.017), membrane (magnitude/breadth, p<0.001/p=0.045), ORF3 (magnitude/breadth, p<0.001/0.001) and ORF8 (magnitude/breadth, p=0.011/0.026) proteins (Figure 2).

**Figure 1:**
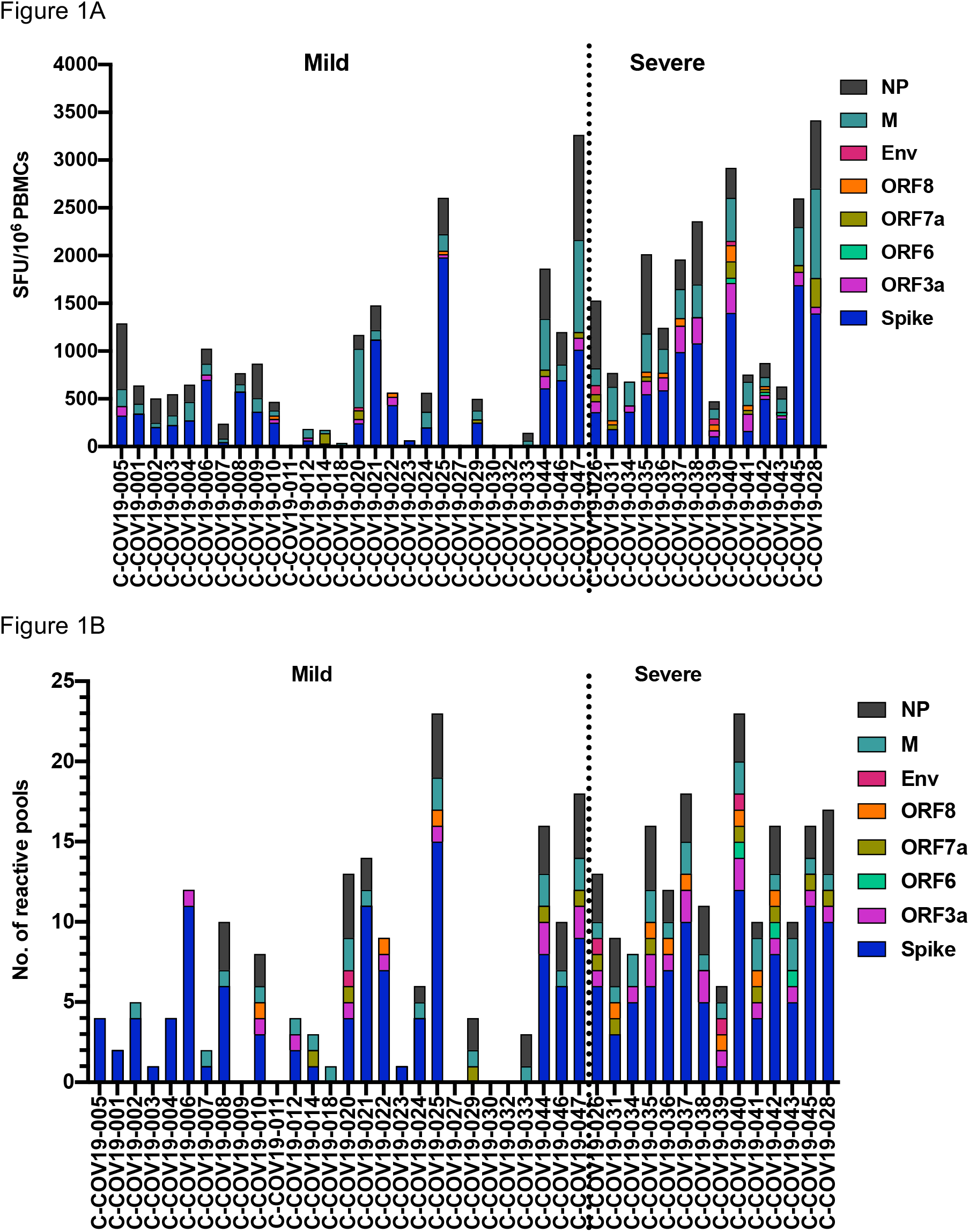
Memory T cell responses specific to SARS-CoV-2 virus proteins in 42 convalescent SARS-CoV-2-infected patients. 28 individuals had mild symptoms while 14 showed severe symptoms. PBMC were isolated and IFNγ production was detected by ELISpot after incubation with SARS-CoV-2 peptides. A) Magnitude of IFNγ T cell responses from each individual. Each bar shows the total T cell responses of each individual specific to all the SARS-CoV-2 protein peptides tested. Each colored segment represents the source protein corresponding to peptide pools eliciting IFNγ T cell responses. B) Breadth of T cell responses from each individual. The breadth of T cell responses was calculated by the number of peptide pools cells responded to. SFU spot forming units

**Figure 2:**
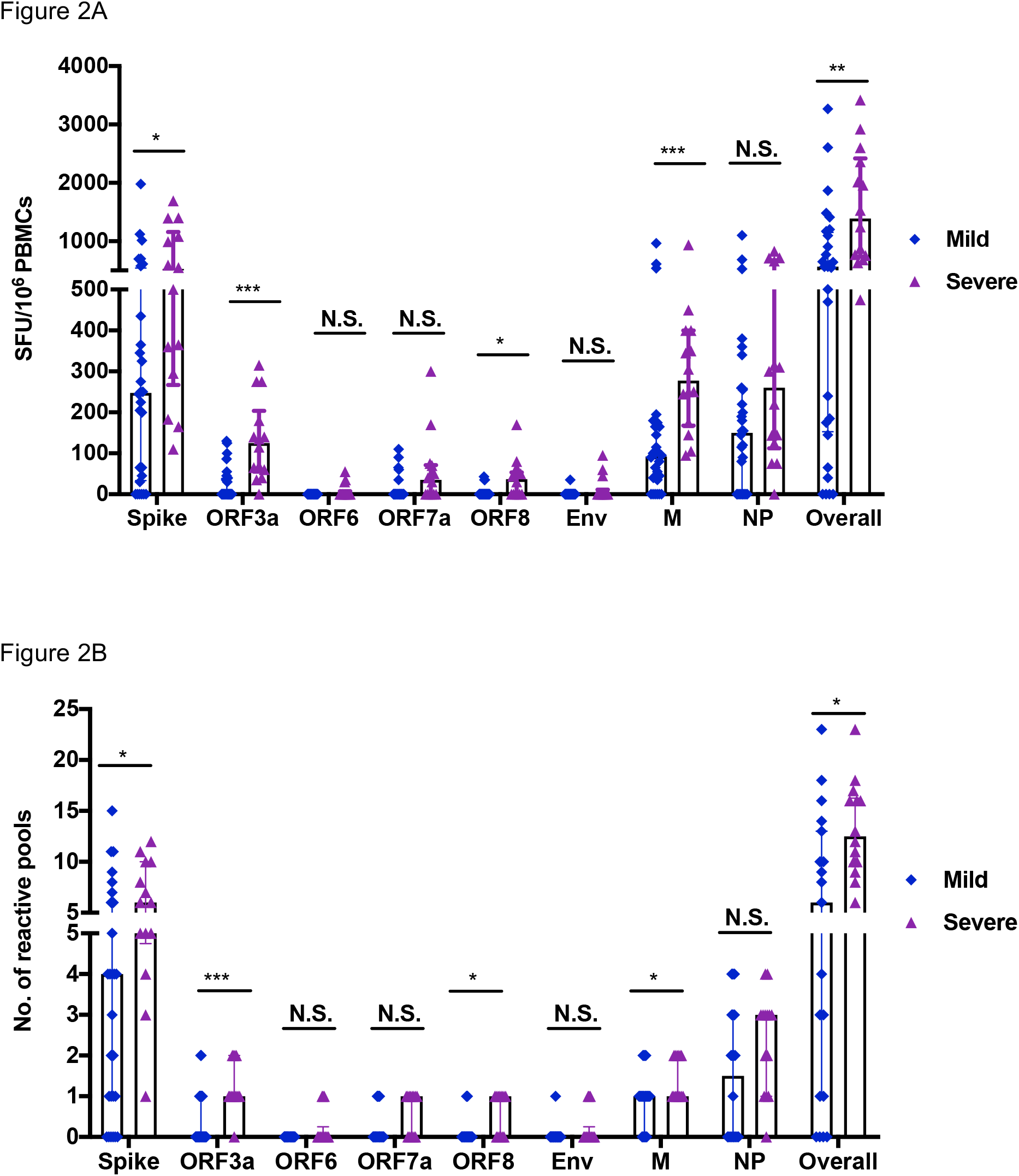
Comparison of magnitude and breadth of T cell response specific to each viral protein between convalescent patients with mild symptoms and severe symptoms. PBMCs were isolated and IFNγ production was detected by ELISpot after incubation with SARS-CoV-2 peptides. A) and B) illustrate the magnitude and the breadth of T cell response against each viral protein between the groups with mild symptoms (n=28) and with severe symptoms (n=14), respectively. SFU spot forming units; Mann-Whitney test was used for the analysis *P<0.05, **P<0.01, ***P<0.001.

### Correlation with spike specific antibody responses

We have assessed the relationship between spike-specific, and overall T cell responses in association with spike-specific, RBD and NP antibody endpoint titres (EPTs). We found significant correlations between (a) spike-specific antibody titers and both overall T cell responses (p<0.001/R=0.52) and spike-specific T cell responses (p=0.001/R= 0.51); (b) RBD-specific antibody titers and both overall T cell responses (p<0.001/R=0.52) and spike-specific T cell responses (p<0.001/R=0.52); and (c) NP-specific antibody titers and both overall T cell responses (p=0.002/R=0.47) and spike-specific T cell responses (p=0.007/R=0.41). However, there was no significant association between NP-specific antibody titers and NP-specific T cell responses (p=0.067/R= 0.29); (Figure 3A-C; and Supplementary Figure 4). Moreover, significantly higher level of spike, RBD and NP EPTs were observed in severe cases in comparison with mild cases (Figure 3D).

**Figure 3:**
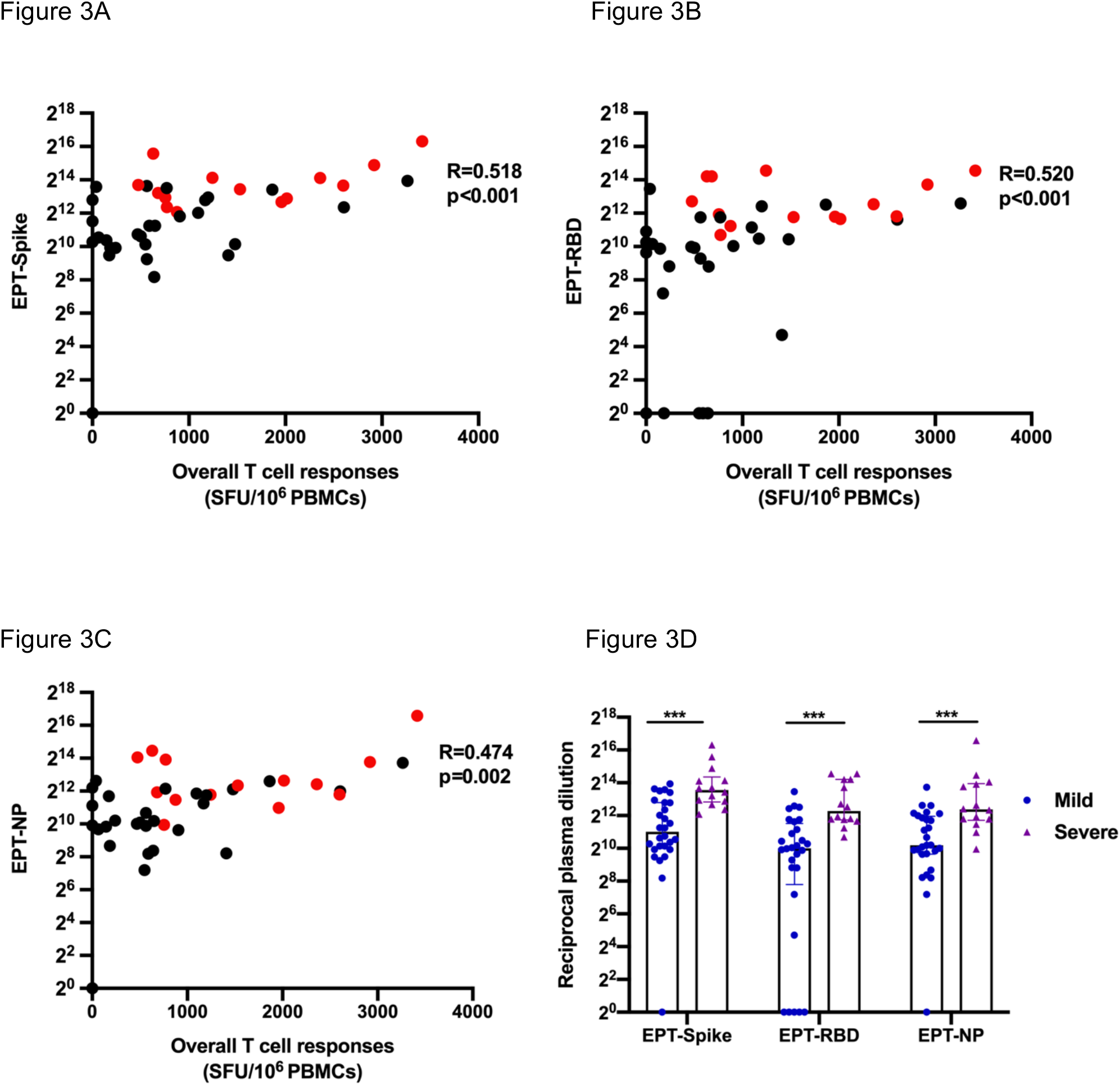
Correlation of T cell responses against SARS-CoV-2 with Spike, RBD and NP-specific antibody responses. A) EPTs-spike B) EPTs-RBD and C) EPTs-NP in association with overall T cell responses. Red dots represent the patients with severe symptoms whereas the mild cases are shown as black dots. D) Comparison of EPT-spike, EPT-RBD and EPT-NP with mild symptoms and severe symptoms. n=42. Spearman’s rank correlation coefficient was used for the correlation analysis and Mann-Whitney test was used for comparison. *** P<0.001; EPT: Endpoint titer

### Distribution of SARS-CoV-2–specific CD4^+^ and CD8^+^ memory T cell responses

Having identified overall T cell responses to SARS-CoV-2 peptides, the responses detected against positive peptide pools were characterized further for involvement of CD4^+^ or CD8^+^ T cell subsets in peptide recognition by evaluating specific intracellular production of IFN-γ and/or TNF-α and/or IL-2 by T cells upon stimulation (Figure 4A, 4B). A greater proportion of the T cell responses detected to spike- and M/NP were found to be mediated by CD8+ (rather than CD4+) T cells in those with mild disease compared to those with severe disease (Figure 4C).

**Figure 4:**
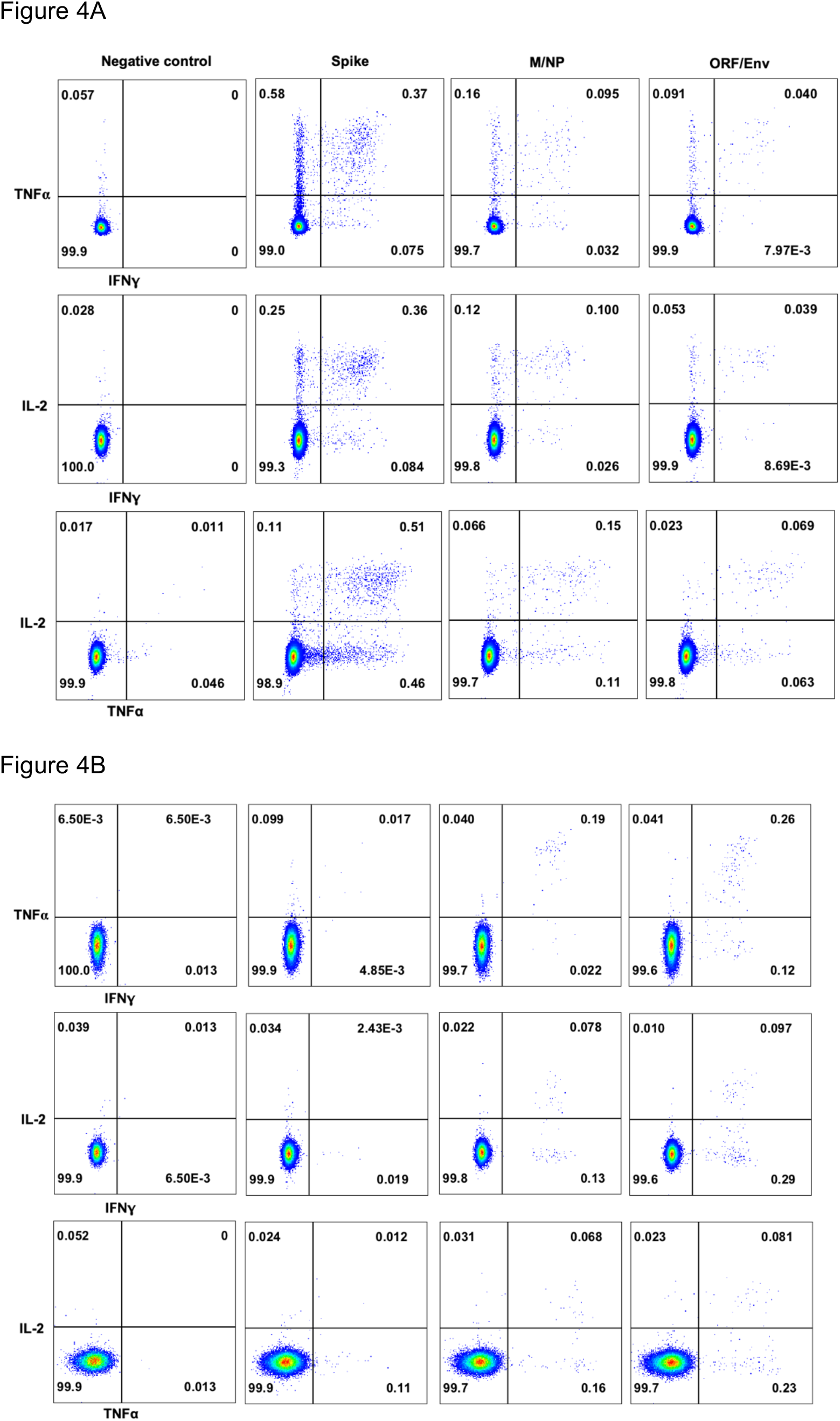
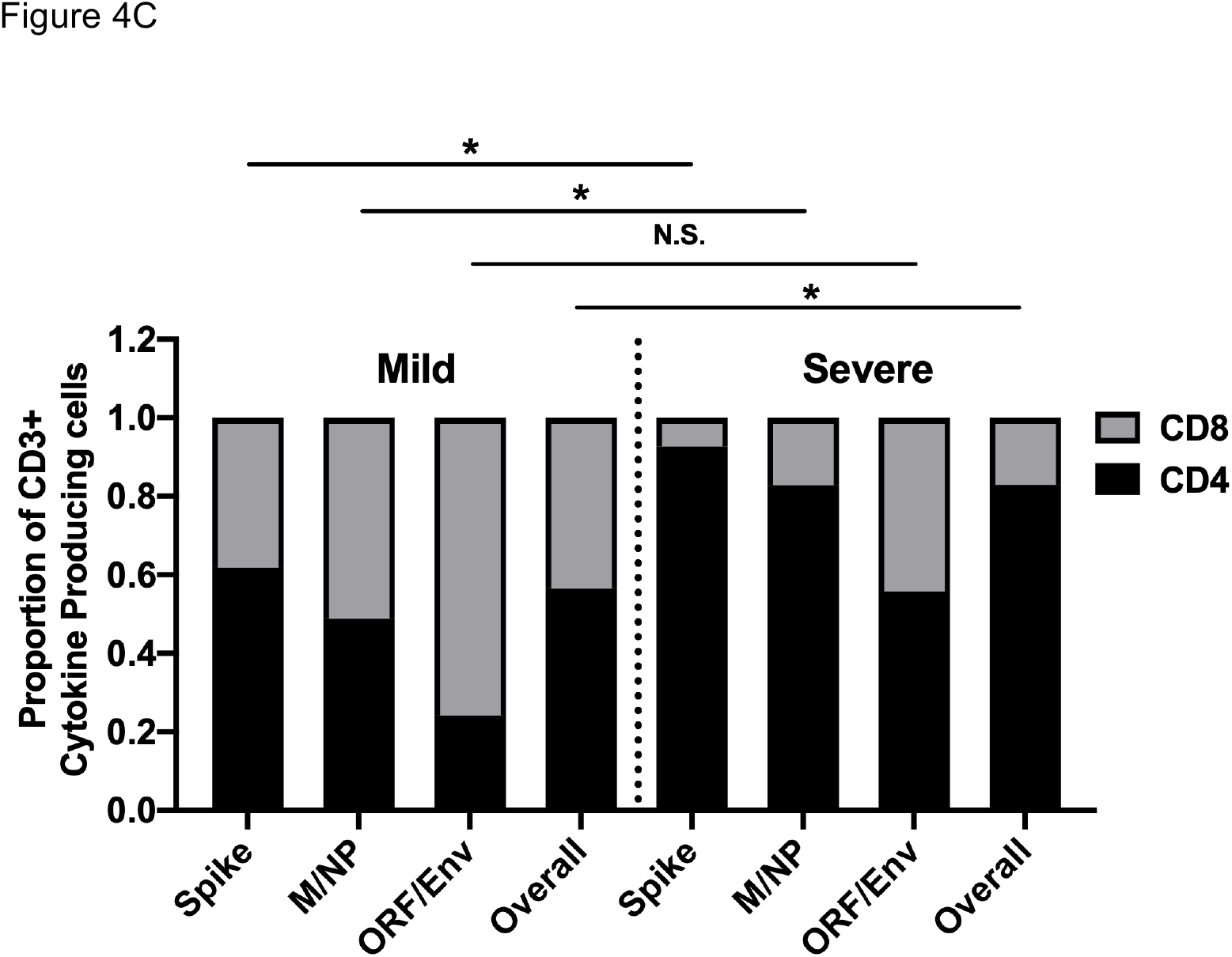
Distribution of SARS-CoV-2–specific CD4^+^ and CD8^+^ memory T cell responses. Cytokine producing T cells were detected by intracellular cytokine staining (ICS) after incubation with SARS-CoV-2 peptides. A) and B) FACS plots representative of CD4^+^T cell and CD8^+^ T cells response respectively upon stimulation with respective SARS-CoV-2 peptide pools. C) The relative proportion of SARS-CoV-2 peptide pool-reactive CD4^+^ and CD8^+^ T cells. The SARS-CoV-2 peptide pool-reactive CD4^+^ or CD8^+^ T cells were identified with at least one of the three cytokines detected: IFN-γ, TNF-α and IL-2. Data shown are from 14 subjects with previously mild COVID-19 symptoms and 8 with severe symptoms. Mann-Whitney test was used for the analysis. * P<0.05

### Evaluation of the polyfunctionality of T cells responding to SARS-CoV-2 peptides

Multi-cytokine analysis revealed a similar pattern of IFNγ, TNFα and IL-2 production by CD4^+^ and CD8^+^ T cells in both mild and severe cases (Figure 5A), For 18 individuals tested, both CD4^+^ and CD8^+^ antigen-specific-T cells produced least one of these three cytokines and others in combination. Higher proportions of CD4^+^T cells were able to produce more than one cytokine than CD8^+^ T cells (Figure 5B). There was no difference in the overall number of multifunctional T cells between the mild group and severe group. Interestingly, CD8^+^ T cells targeting different virus proteins showed different cytokine profiles, with the M/NP-specific CD8^+^ T cells having wider functionality than T cells targeting spike protein (p=0.0231, Figure 5B and 5C). Furthermore, the M/NP-specific CD8^+^ T cells were significantly higher than spike-specific T cells in those with mild disease, but not in those with severe disease. In contrast to observations seen in influenza virus infection^21^, we did not observe significant differences in the cytotoxic potential (as indicated by expression of the degranulation marker CD107a) in patients with mild and severe disease (Figure 5D); and we observed very few CD107a^+^ CD4^+^ T cells overall, suggesting cytotoxic CD4^+^ T cells might not be a major contributor to virus clearance.

**Figure 5:**
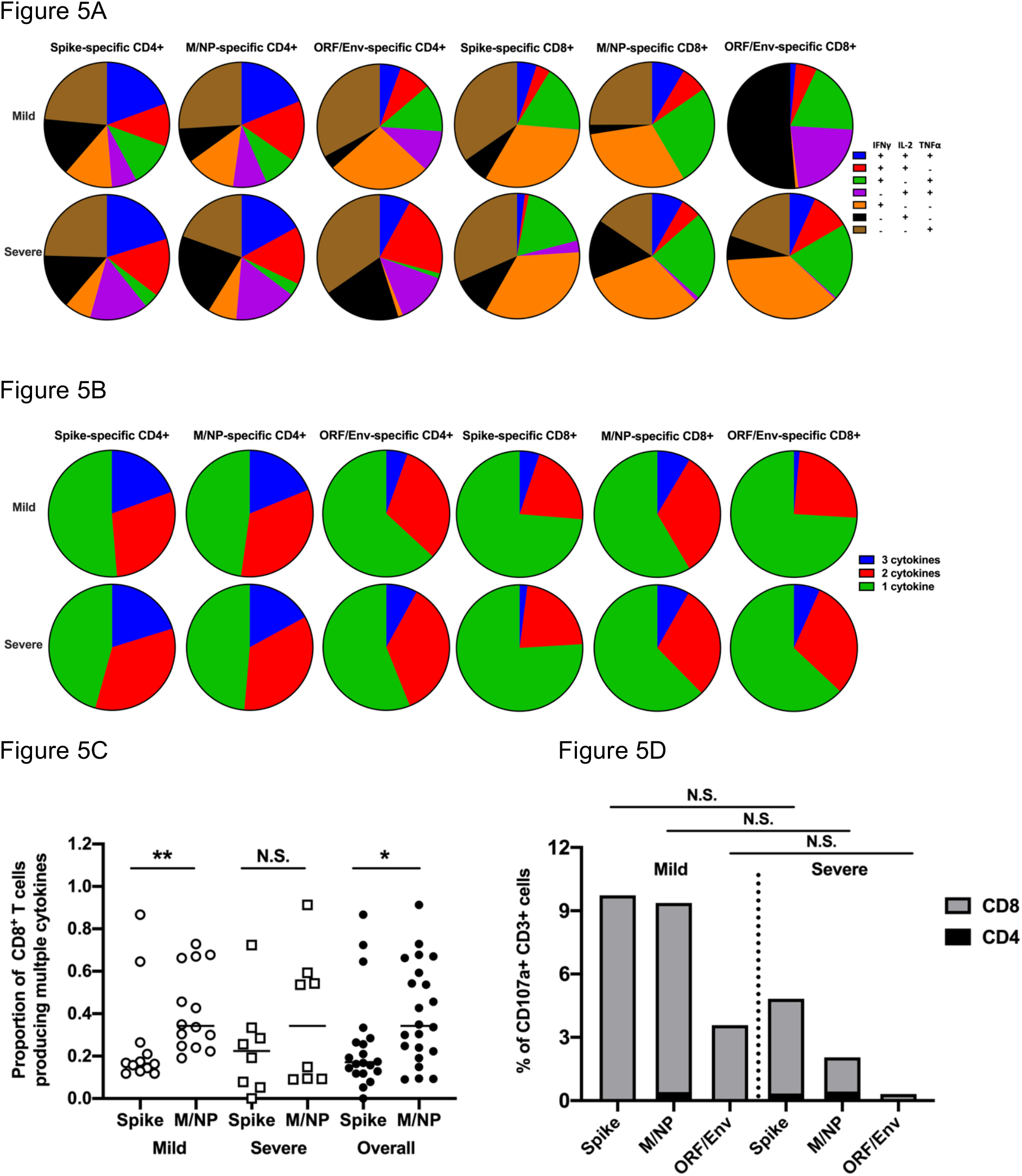
Cytokine profile of SARS-Cov-2-specific T cells. Cytokine production of SARS-Cov-2-specific T cells was assessed by intracellular cytokine staining after incubation with SARS-CoV-2 peptides. Pie charts (A) and (B) represent the relative proportions of CD4^+^ or CD8^+^ T cells producing IFN-γ and/or TNFα, and/or IL-2, and the relative proportion of T cells producing one, two and three cytokines, respectively. Different colored segments represented different pattern of cytokine production. C) Comparison of the frequency of multifunctional CD8^+^ T cells targeting Spike and M/NP. The open circles and squares represent T cell responses in patients with mild symptoms and the ones with severe symptoms, respectively. D) The relative frequencies of CD4^+^ and CD8^+^ T cells expressing CD107a after antigen-stimulation. Data shown are from 14 subjects with mild symptoms and 8 with severe symptoms. Mann-Whitney test was used for the analysis. * P<0.05, **P<0.01

### Identification of SARS-CoV-2 specific T cell peptides containing epitopes

IFNγ ELISpot assays were performed with candidate peptides identified from the 2-dimensional matrix analysis in 34 subjects. A total of 39 peptides containing SARS-CoV-2 T cell epitope regions were recognized by COVID-19 recovered subjects, 17 from spike, 10 from NP, 6 from membrane and 7 from ORF proteins. Strikingly, 6 dominant peptides were recognised by more than 18% of subjects tested (Table 1). NP-16 was recognised by 12/34 (35%) subjects tested and contained at least two epitopes which were CD8^+^ or CD4^+^ dependant.

**Table 1.**
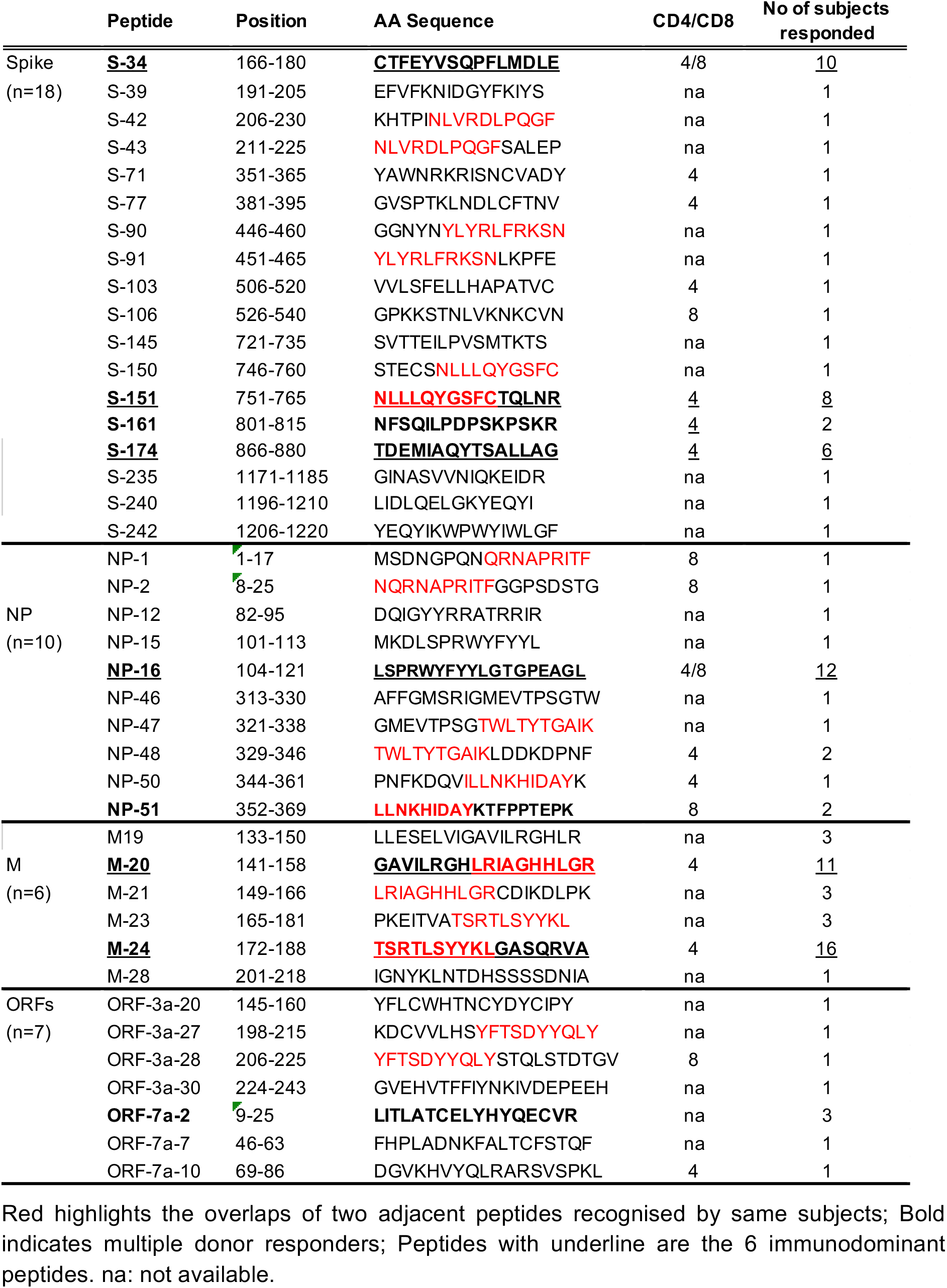
Peptides containing T cell epitopes.

M-24 was recognised by 16/34 subjects (47%) tested and contained one or more CD4^+^ epitopes. Peptide M-20 was recognised by 11/34 subjects tested (32%) and contained one or more CD4^+^ epitopes. 3 dominant spike peptides were also identified, with S-34 recognised by 10/34 subjects (29%) containing both CD4^+^ and CD8^+^ epitopes, and a further two spike peptides S-151 and S-174 were recognised by 8/34 and 6/34 subjects (24% and 18%), both containing CD4^+^ epitopes.

Those dominant responses were further confirmed by both *ex-vivo* and using cultured short-term T cell lines. Supplementary Figure 5 illustrates example FACS plots from intracellular cytokine staining (ICS) when short-term T cell lines were stimulated with single peptides containing epitopes. CD4^+^ T cells elicited strong responses against dominant spike peptides and M peptides, whereas cells targeting two NP dominant peptides were CD8^+^ T cells. The optimal epitopes recognised within long peptides to which dominant CD8+ responses were detected and their HLA restriction were predicted using the optimal epitope prediction run by IEDB analysis tool (http://tools.iedb.org/mhci). The best predicted epitope sequences identified are shown in Table 2.

**Table 2:**
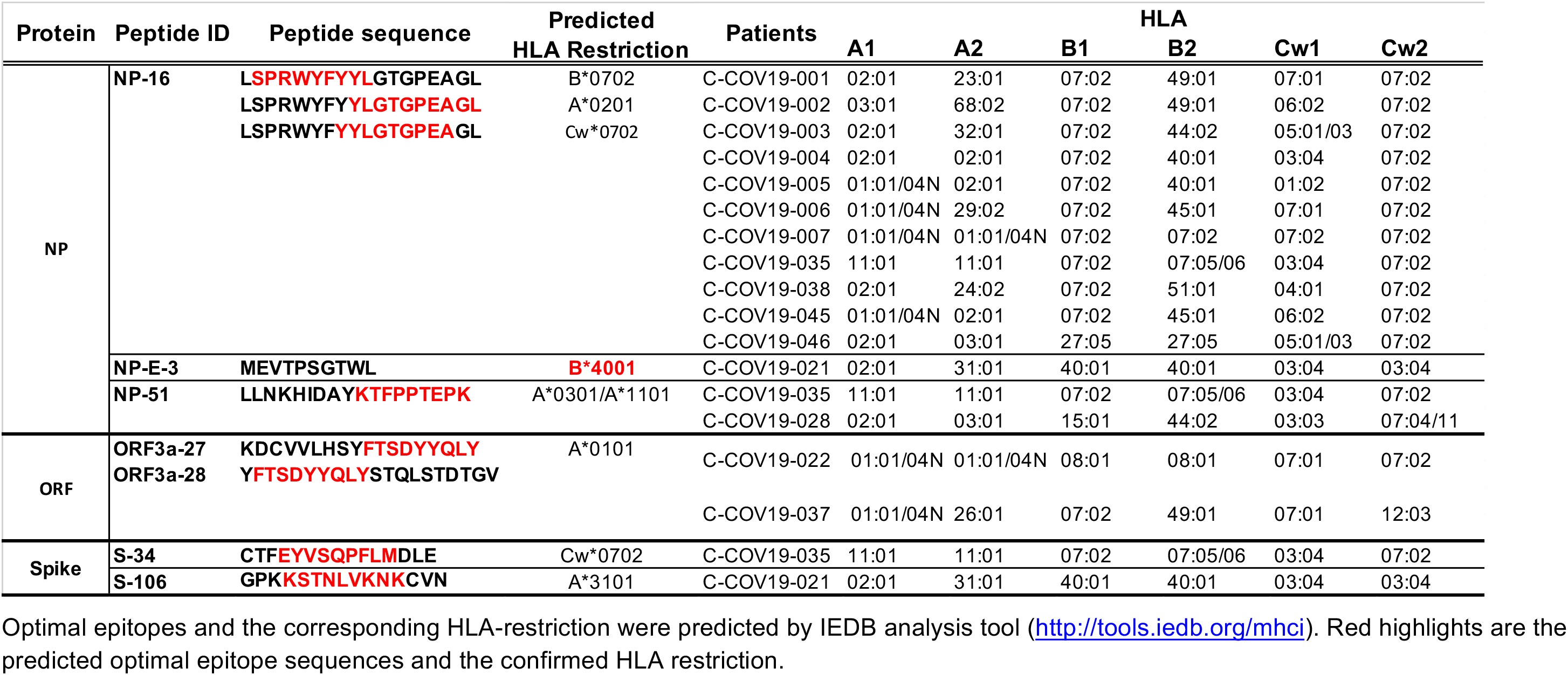
HLA class I typing of CD8^+^ epitope peptides in subjects with confirmed responses.

A set of previously defined SARS epitopes^16^ with identical sequences to SARS-CoV-2 were also tested by ELISpot assay (Supplementary table 2), Most of those peptides did not elicit any positive responses in 42 COVID-19 recovered subjects, apart from two NP epitope peptides (N-E-3 MEVTPSGTWL and N-E-11 LLNKHIDAYKTFPPTEPK) and one spike epitope peptide (S-E-19 QLIRAAEIRASANLAATK). The HLA restriction of epitope N-E-3 was confirmed by flow cytometry analysis of HLA Pentamer complexes of N-E-3 to be HLA-B*4001. The majority of N-E-3-specific CD8^+^ T cells exhibited central memory (26.8%) or effector memory phenotype (54.6%) (Figure 6) and early (CD27+CD28+, 56%) or intermediate (CD27+CD28−, 43.3%) differentiation phenotype. In addition to this well-defined NP epitope, N-E-11, which is identical to peptide NP-51, shares the sequence with two other known HLA-A*0201 restricted SARS epitopes (N-E-1 ILLNKHID and N-E-5 ILLNKHIDA). Interestingly, one of the responders to this peptide did not carry the HLA-A*0201 allele (Table 1), indicating this peptide may contain a different SARS-CoV-2 epitope presented by a different HLA molecule. Whereas these NP epitopes are targeted by CD8^+^ T cells, we also detected a CD4^+^ T cell response targeting SARS spike epitope S-E-19 which spans between the overlapping peptides of S-203 and S-204. This peptide is known to be presented by HLA-DRB1*0401 in SARS infection.

**Figure 6:**
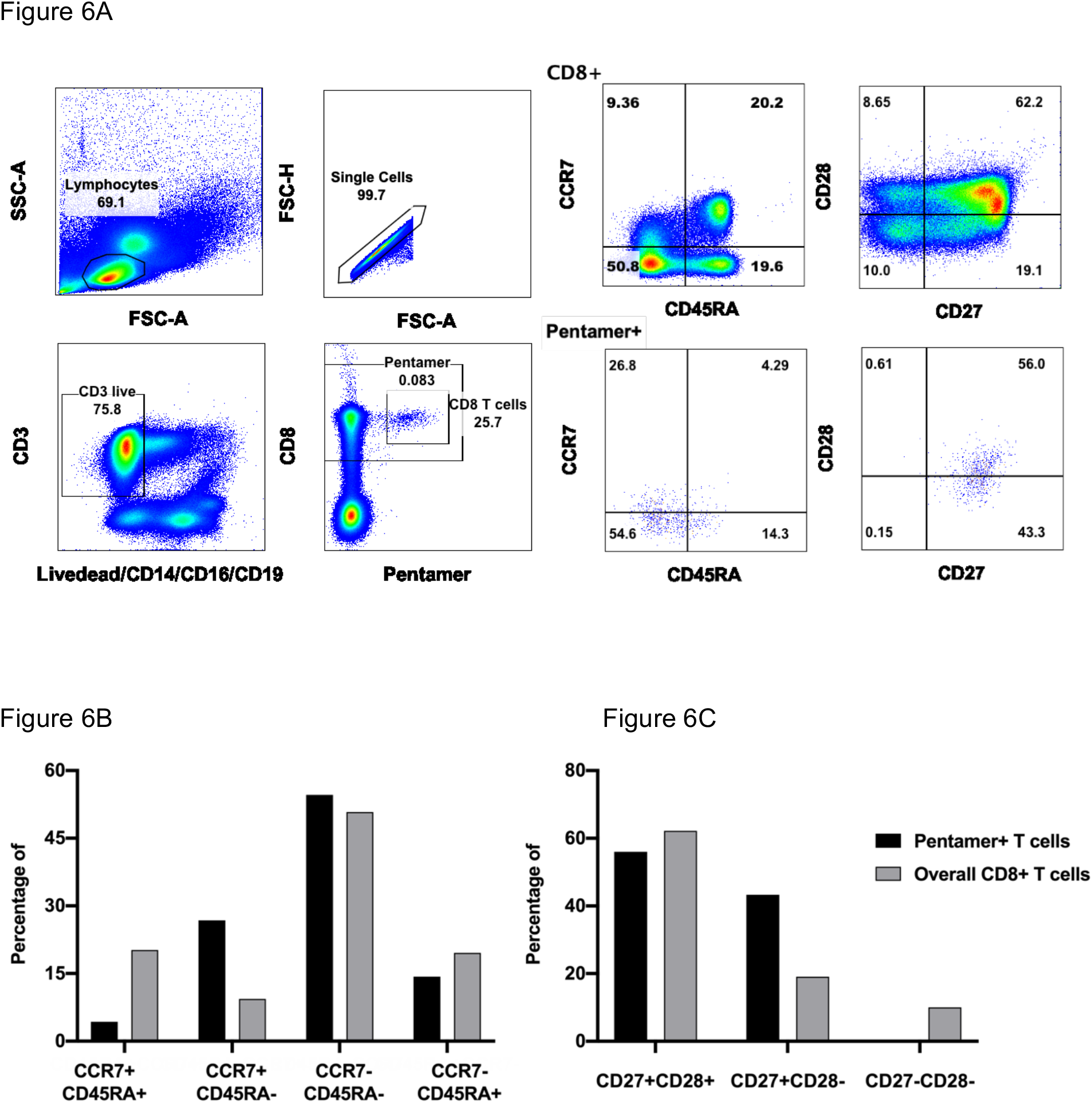
Memory phenotype and differentiation status of HLA-B*4001 restricted NP_322-331_ (MEVTPSGTWL)-specific T cells. PBMC were isolated and stained with NP/B*4001 Pentameric complexes and markers of T cell differentiation, and analysed using flow cytometry. A) Expression of differentiation markers of CCR7 and CD45RA on CD8^+^ Pentamer+ T cells. B): Expression of differentiation markers of CD27 and CD28 on CD8^+^ Pentamer+ T cells.

Pre-existing cross-reactive T cell responses to common coronaviruses might play a role in immunity to COVID-19^15^. However, we found the alignment of all epitopes identified here, showed little homology with other common coronaviruses (Supplementary Figure 6).

## Discussion

This is the first in-depth analysis of SARS-CoV-2 specific T cell responses in individuals who have recovered from COVID-19 stratified for disease severity. We found the breadth and magnitude of the memory T cell responses from convalescent individuals with COVID-19 to be greater in severe compared to mild cases, and this was most markedly observed to spike, M, and ORF3 proteins. M/Np-specific CD8^+^ T cells were significantly higher frequency than spike-specific CD8^+^ T cells in those with mild disease. The overall T cell responses as well as spike-specific T cell responses correlated with spike, RBD and NP-specific antibody end point titre. We also identified 39 individual peptides containing CD4^+^ and/or CD8^+^ epitopes with 6 immunodominant epitope clusters defined. The memory responses we identified were fully functional and capable of producing IFN-γ TNF-α and IL-2. CD8^+^ T cells expressed the degranulation marker CD107a consistent with the killing potential of CD8^+^ cytotoxic T cells. Furthermore, the ratio of CD8:CD4 SARS-CoV-2-specific T cells was greater for those with a history of mild disease. Our data highlight the potential importance of including non-spike proteins in future COVID-19 vaccine design.

Overall broader and stronger SARS-CoV-2 specific T cell responses in patients who recovered from severe disease may reflect the higher viral loads, or an insufficient response to elicit early control of virus or that the response was itself deleterious. Consistent with the Grifoni *et al* recent report ^15^, we also observed a particularly high frequency of spike protein-specific CD4^+^ T cell responses in patients who had recovered from COVID-19. This is very similar to influenza virus infection, where viral surface hemagglutinin (HA) expressed on the surface of influenza virus elicited mostly CD4^+^ responses, whereas the majority of CD8^+^ responses were specific to viral internal proteins ^17^. Understanding the immune protection by different subsets of T cells in mild cases and their potential roles played in COVID-19 pathogenesis is a crucial question. The timing and strength of early primed T cell responses, could be critical determinants of viral control at an early stage of the infection.

Among the total 39 peptides containing T cell epitopes identified in this study, strikingly, we found six immunodominant epitope clusters (peptides) frequently targeted by T cells in many donors, including 3 clusters in spike (29%, 24%, 18%), two in membrane protein (32%, 47%) and one in nucleoprotein (35%). The high frequency of immunodominant responses observed could be due to high level of virus co-receptor ACE2 expression ^22^ and high binding affinity to spike of SARS-CoV-2 ^23^ allowing more viral entry and more viral protein available for antigen presentation. Whether or not these dominant responses play a role in immune protection merits further investigation in a larger cohort, especially to include those who are symptomatic with COVID-19.

We observed multiple strong dominant CD8^+^ T cell responses in study subjects specific to the NP protein with dominant epitope clusters within NP (NP-16) detected in 35% study subjects. In addition, a higher frequency of multi-cytokine producing M/Np-specific than spike-specific CD8^+^ T cells was observed in subjects who had recovered from mild disease. These findings suggest Np and M have potential as candidate proteins for inclusion within future vaccine design.

However, we did not observe frequent IFN-γ producing SARS-CoV-2 specific T cell responses in healthy unexposed volunteers which is different from recently published data by Grifoni *et al*^15^ which was also observed by Braun *et al* ^24^. Our data are supported by a recent immunogenicity study of a recombinant adenovirus type-5 (Ad5) vectored COVID-19 vaccine human phase I trial (108 volunteers without pre-exposure to COVID-19), where spike-specific T cell responses before vaccination were not found when using IFN-γ based assays ^7^. We believe the differences could be caused by differences in sensitivity of detection method used between the studies, where Grifoni *et al* have used Activation Induced Marker assays (AIM) as their main detection method for antigen-specific T cells. IFN-γ based detection assays (ELISpot and intracellular cytokine staining) are well-established methods to evaluate antigen specific T cells in different virus infections, with direct functional relevance ^17, 25, 26, 27^, The AIM assay is more recently developed assay, capable of detecting responsive T cells independently of individual cytokines. Alternatively, it is possible that different circulating coronaviruses have been recently been present in the different populations studied.

The dominant epitopes identified in this study showed little similarity between SARS-CoV-2 and other coronaviruses raising the question of the level and impact of pre-existing cross-reactive T cells contributing to the epitopes that we identified. This is very different from what we observed in influenza virus infection during the H5N1 pandemic where 90% homology was observed between different strains with a high level of cross-reactivity detected by interferon-gamma based assays ^17^. The presence and functionality of T cells in healthy volunteers without previous exposure to COVID-19, requires further investigation in different populations using different approaches.

Taken together, our study has demonstrated strong and broad SARS-CoV-2-specific T cell responses in the majority of subjects who had recovered from COVID-19. Immunodominant epitope clusters and peptides containing T cell epitopes identified in this study will provide critical tools to study the contribution of SARS-CoV-19 specific T cells in protection and immune pathology. Identification of non-spike dominant CD8^+^ T cell epitopes suggest the potential importance of including of non-spike protein such as NP, M and ORFs into future vaccine designs.

## Acknowledgments

We are grateful to all of the participants for donating their samples and data for these analyses, and the research teams involved in the consenting, recruitment and sampling of these participants. We acknowledge the support of ISARIC4C Investigators.

This work is supported by UK Medical Research Council; Chinese Academy of Medical Sciences (CAMS) Innovation Fund for Medical Sciences (CIFMS), China (grant number: 2018-I2M-2-002); The National Institute for Health Research [award CO-CIN-01]; the Medical Research Council [grant MC_PC_19059]; the National Institute for Health Research Health Protection Research Unit (NIHR HPRU) in Emerging and Zoonotic Infections at University of Liverpool in partnership with Public Health England (PHE)[NIHR award 200907]; Wellcome Trust and Department for International Development [215091/Z/18/Z]; the Bill and Melinda Gates Foundation [OPP1209135]; NIHR Oxford Biomedical Research Centre for support and Centre and National Institutes of Health, National Institute of Allergy and Infectious Disease (Consortium for HIV/AIDS Vaccine Development UM1 AI 144371 (PB and AMcM) and R01 AI 118549 (PB));

The views expressed are those of the authors and not necessarily those of the DHSC, DID, NIHR, MRC, Wellcome Trust or PHE.

---

**Supplementary Figure 1.**
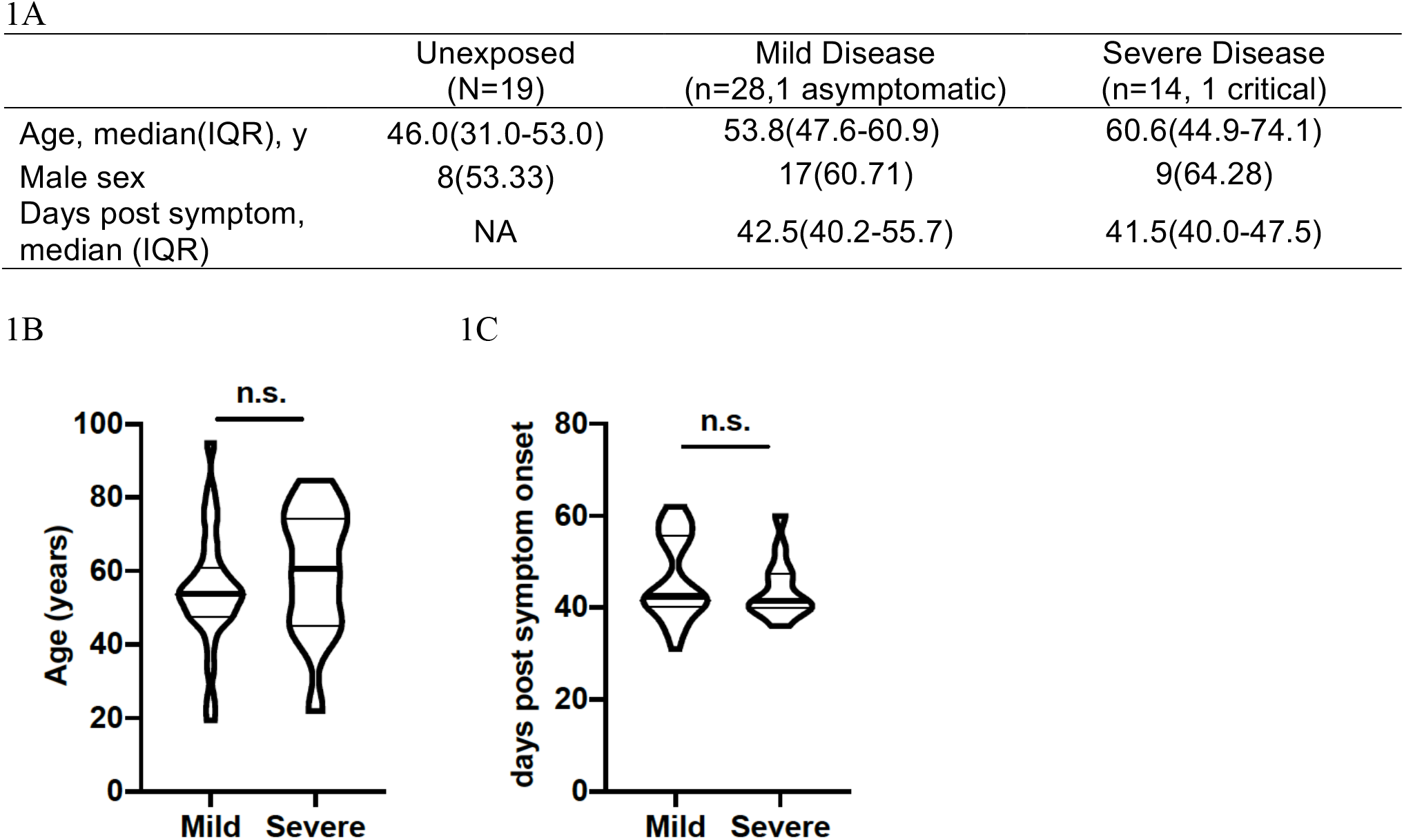
Participant characteristics.

**Supplementary Figure 2.**
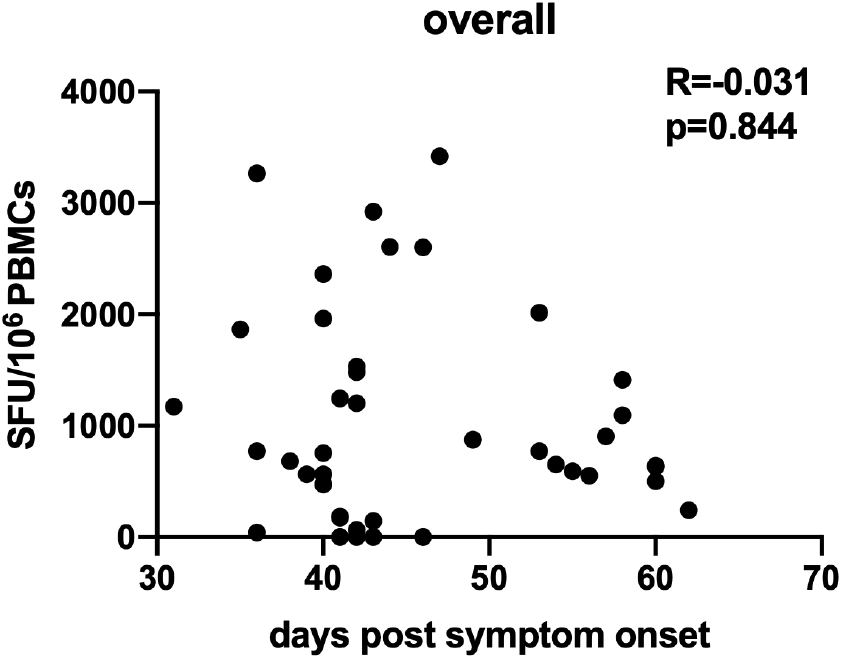
Correlation between overall T cell response of each individual and the days post symptom when blood specimen was taken. n=42

**Supplementary Figure 3.**
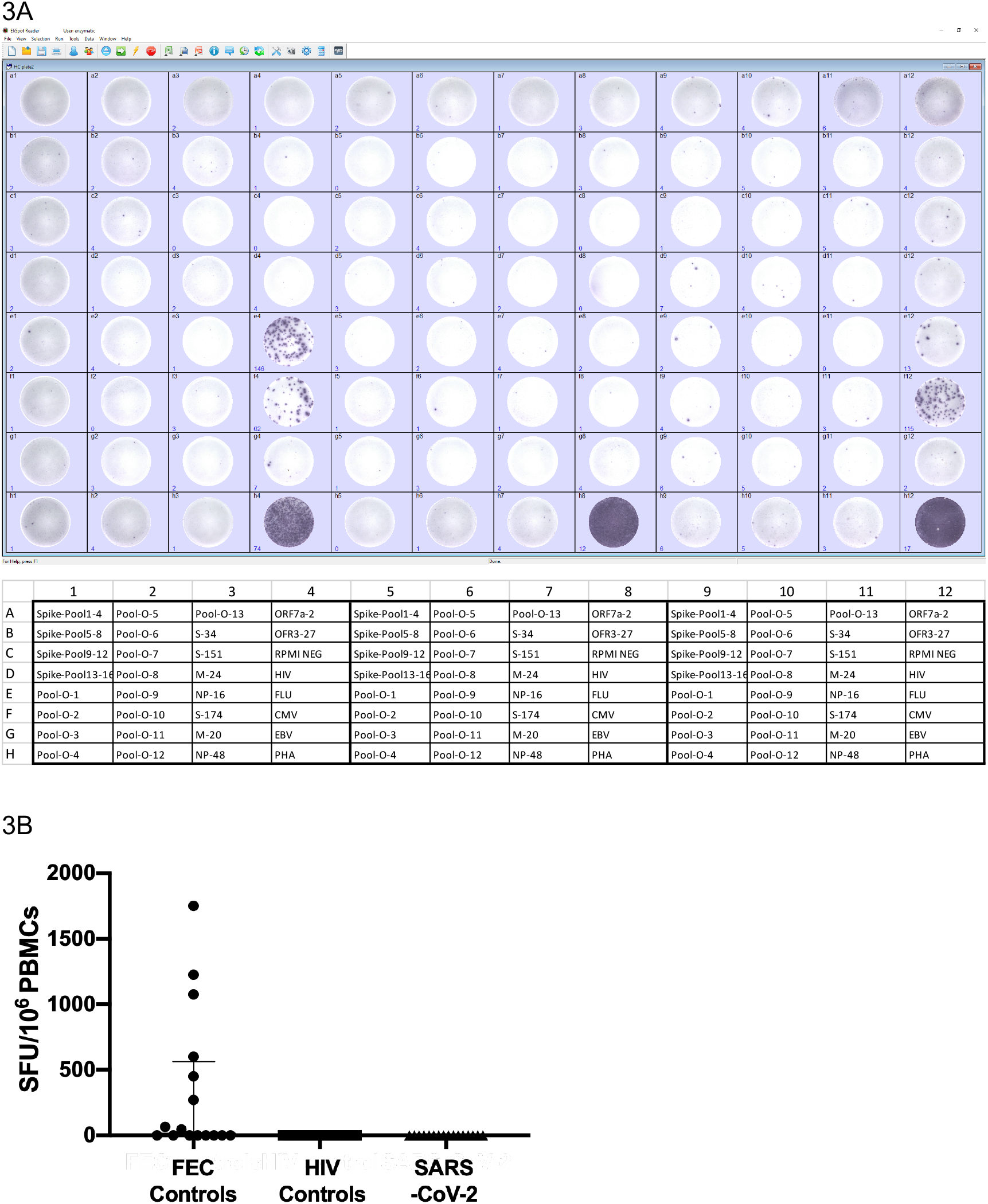
Magnitude of T cell responses of unexposed healthy individuals against SARS-CoV-2 antigens, known CD8^+^ T cell epitopes of human influenza, CMV and EBV viruses (namely FEC controls) and a peptide pool of irrelevant antigen HIV Gag protein (n=15). A) An Example of IFN-γ ELISpot plate from three healthy individuals without SARS-Cov-2 infection. Each individual has been tested with four big spike pools (Pool 1-4, Pool 5-8, Pool 9-12 and Pool-13-16), 13 first dimension of non-spike pools and nine dominant individual peptides containing epitope, along with six control wells. B). Magnitude of T cell responses of 15 unexposed healthy individuals against SARS-CoV-2 antigens and control antigens

**Supplementary Figure 4.**
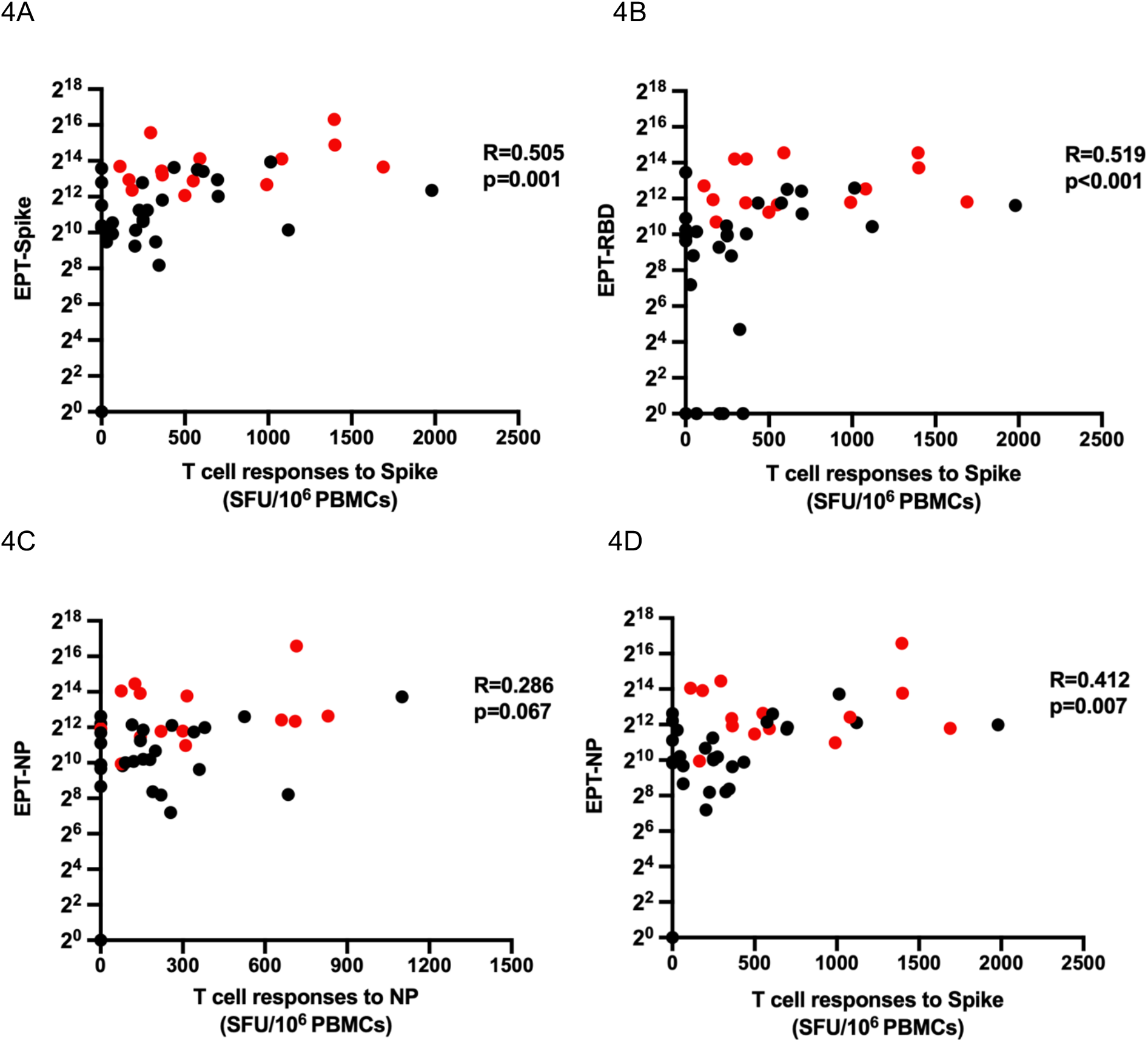
Correlation of Spike, RBD and NP-specific T cell responses with corresponding SARS-CoV-2 antigen-specific antibody responses. n=42

**Supplementary Figure 5.**
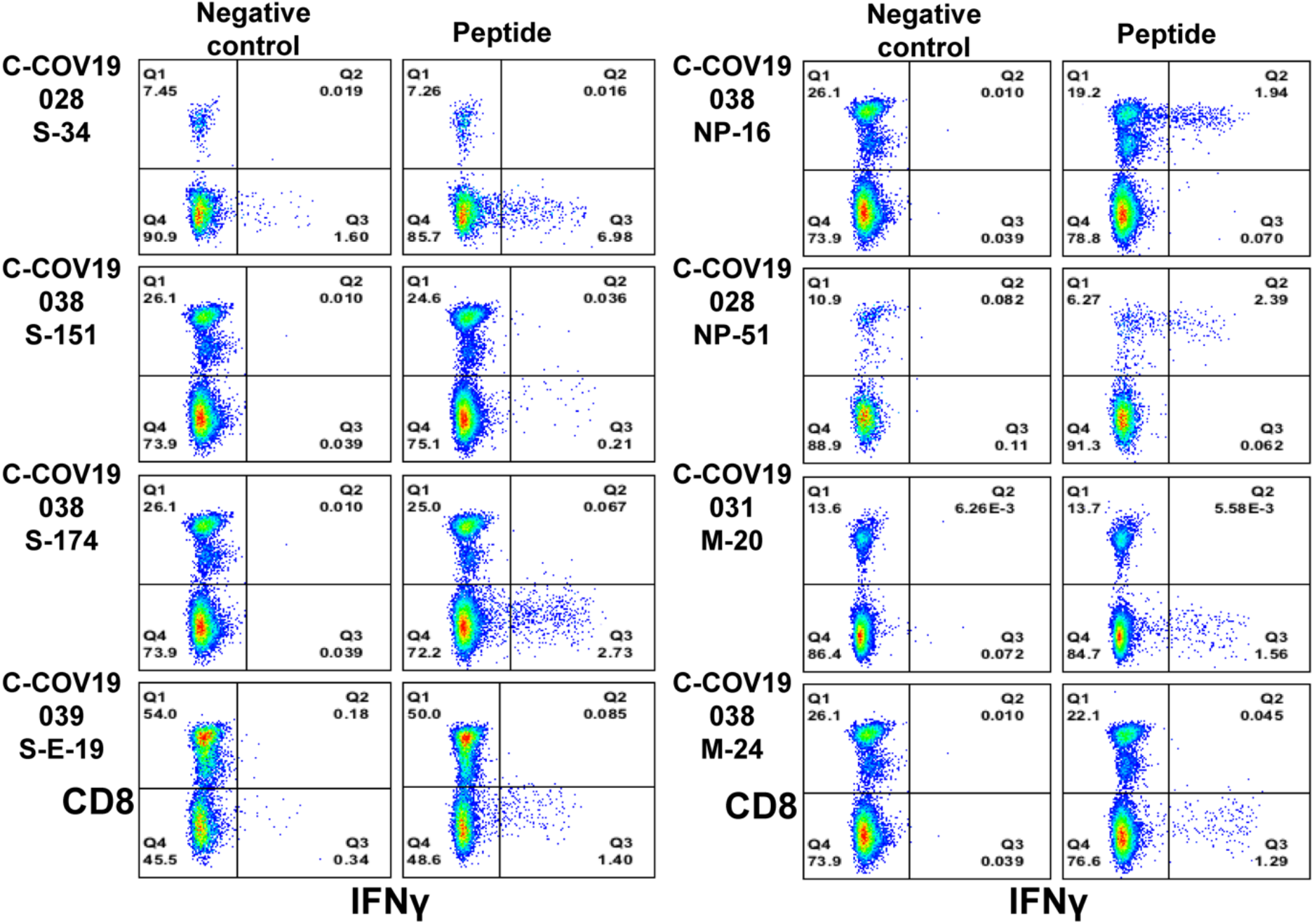
Confirmation of dominant T cell response with cultured short-term T cell lines. PBMCs were stimulated with corresponding peptide pools according to the ex vivo ELISpot results and then cultured for 10 days. Cytokine production of the cell lines was then examined by ICS upon the stimulation with single peptides.

**Supplementary Figure 6:**
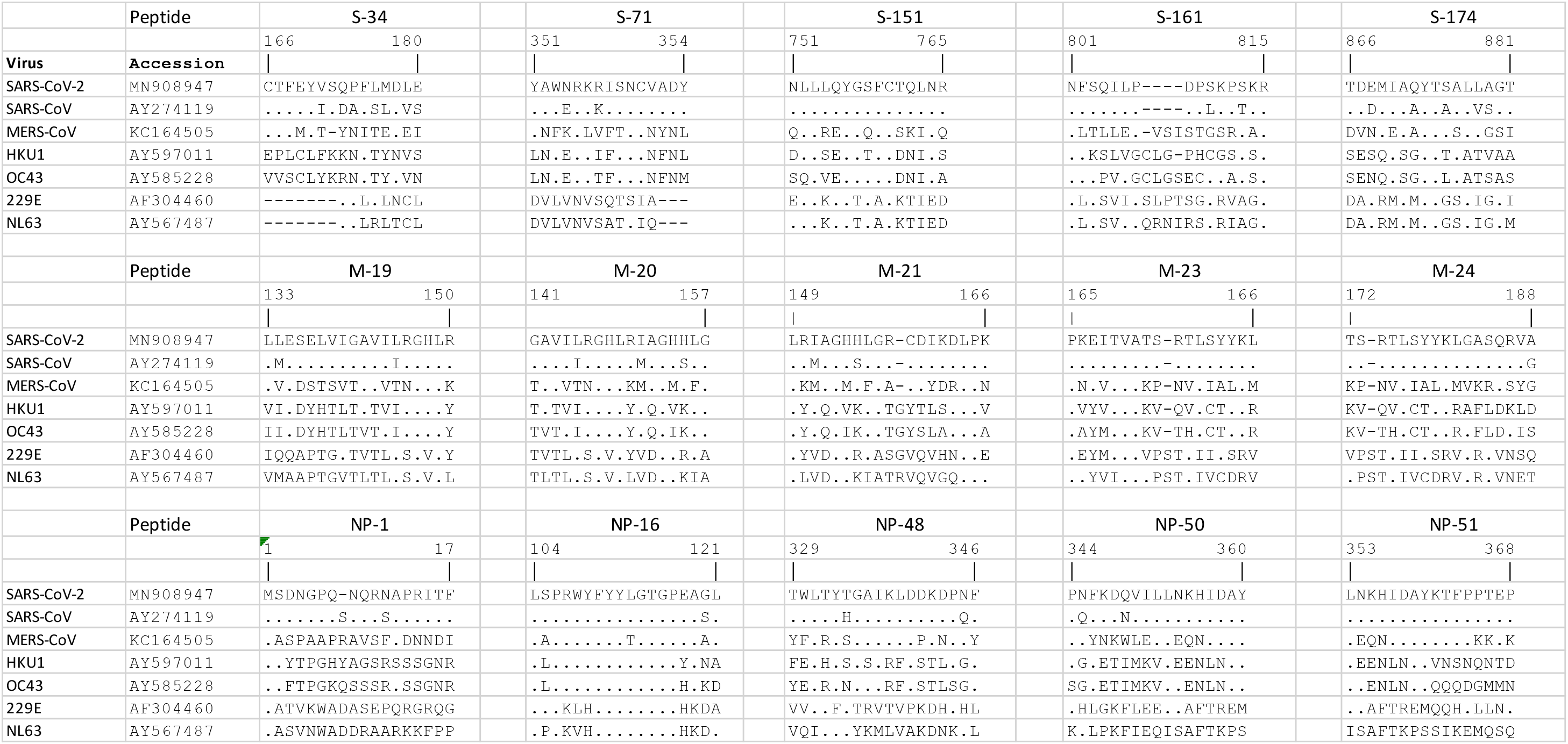
Alignments of dominant peptides identified with common coronaviruses. “-” represents a gap between the amino acids, “.” stands for the identical amino acid.

**Supplementary Table 1:** Two-dimensional peptide Matrix pools of spike protein (A) and non-spike proteins including ORF3a, OFR6, ORF7a, ORF8, Envelope, Membrane Protein and Nucleoprotein (B).

|  | Pool-17 | Pool-18 | Pool-19 | Pool-20 | Pool-21 | Pool-22 | Pool-23 | Pool-24 | Pool-25 | Pool-26 | Pool-27 | Pool-28 | Pool-29 | Pool-30 | Pool-31 | Pool-32 |
| --- | --- | --- | --- | --- | --- | --- | --- | --- | --- | --- | --- | --- | --- | --- | --- | --- |
| Pool 1 | S-1 | S-2 | S-3 | S-4 | S-5 | S-6 | S-7 | S-8 | S-9 | S-10 | S-11 | S-12 | S-13 | S-14 | S-15 | S-16 |
| Pool 2 | S-17 | S-18 | S-19 | S-20 | S-21 | S-22 | S-23 | S-24 | S-25 | S-26 | S-27 | S-28 | S-29 | S-30 | S-31 | S-32 |
| Pool 3 | S-33 | S-34 | S-35 | S-36 | S-37 | S-38 | S-39 | S-40 | S-41 | S-42 | S-43 | S-44 | S-45 | S-46 | S-47 | S-48 |
| Pool 4 | S-49 | S-50 | S-51 | S-52 | S-53 | S-54 | S-55 | S-56 | S-57 | S-58 | S-59 | S-60 | S-61 | S-62 | S-63 | S-64 |
| Pool 5 | S-65 | S-66 | S-67 | S-68 | S-69 | S-70 | S-71 | S-72 | S-73 | S-74 | S-75 | S-76 | S-77 | S-78 | S-79 | S-80 |
| Pool 6 | S-81 | S-82 | S-83 | S-84 | S-85 | S-86 | S-87 | S-88 | S-89 | S-90 | S-91 | S-92 | S-93 | S-94 | S-95 | S-96 |
| Pool 7 | S-97 | S-98 | S-99 | S-100 | S-101 | S-102 | S-103 | S-104 | S-105 | S-106 | S-107 | S-108 | S-109 | S-110 | S-111 | S-112 |
| Pool 8 | S-113 | S-114 | S-115 | S-116 | S-117 | S-118 | S-119 | S-120 | S-121 | S-122 | S-123 | S-124 | S-125 | S-126 | S-127 | S-128 |
| Pool 9 | S-129 | S-130 | S-131 | S-132 | S-133 | S-134 | S-135 | S-136 | S-137 | S-138 | S-139 | S-140 | S-141 | S-142 | S-143 | S-144 |
| Pool 10 | S-145 | S-146 | S-147 | S-148 | S-149 | S-150 | S-151 | S-152 | S-153 | S-154 | S-155 | S-156 | S-157 | S-158 | S-159 | S-160 |
| Pool 11 | S-161 | S-162 | S-163 | S-164 | S-165 | S-166 | S-167 | S-168 | S-169 | S-170 | S-171 | S-172 | S-173 | S-174 | S-175 | S-176 |
| Pool 12 | S-177 | S-178 | S-179 | S-180 | S-181 | S-182 | S-183 | S-184 | S-185 | S-186 | S-187 | S-188 | S-189 | S-190 | S-191 | S-192 |
| Pool 13 | S-193 | S-194 | S-195 | S-196 | S-197 | S-198 | S-199 | S-200 | S-201 | S-202 | S-203 | S-204 | S-205 | S-206 | S-207 | S-208 |
| Pool 14 | S-209 | S-210 | S-211 | S-212 | S-213 | S-214 | S-215 | S-216 | S-217 | S-218 | S-219 | S-220 | S-221 | S-222 | S-223 | S-224 |
| Pool 15 | S-225 | S-226 | S-227 | S-228 | S-229 | S-230 | S-231 | S-232 | S-233 | S-234 | S-235 | S-236 | S-237 | S-238 | S-239 | S-240 |
| Pool 16 | S-241 | S-242 | S-243 | S-244 | S-245 | S-246 | S-247 | S-248 | S-249 | S-250 | S-251 | S-252 | S-253 |  |  |  |

1B
|  | Pool-O-14 | Pool-O-15 | Pool-O-16 | Pool-O-17 | Pool-O-18 | Pool-O-19 | Pool-O-20 | Pool-O-21 | Pool-O-22 | Pool-O-23 | Pool-O-24 | Pool-O-25 | Pool-O-26 | Pool-O-27 | Pool-O-28 | Pool-O-29 |
| --- | --- | --- | --- | --- | --- | --- | --- | --- | --- | --- | --- | --- | --- | --- | --- | --- |
| Pool-O-1 | ORF3a-1 | ORF3a-2 | ORF3a-3 | ORF3a-4 | ORF3a-5 | ORF3a-6 | ORF3a-7 | ORF3a-8 | ORF3a-9 | ORF3a-10 | ORF3a-11 | ORF3a-12 | ORF3a-13 | ORF3a-14 | ORF3a-15 | ORF3a-16 |
| Pool-O-2 | ORF3a-17 | ORF3a-18 | ORF3a-19 | ORF3a-20 | ORF3a-21 | ORF3a-22 | ORF3a-23 | ORF3a-24 | ORF3a-25 | ORF3a-26 | ORF3a-27 | ORF3a-28 | ORF3a-29 | ORF3a-30 | ORF3a-31 | ORF3a-32 |
| Pool-O-3 | ORF3a-33 | ORF3a-34 | ORF3a-35 |  |  |  |  |  |  |  |  |  |  |  |  |  |
| Pool-O-4 | ORF6-1 | ORF6-2 | ORF6-3 | ORF6-4 | ORF6-5 | ORF6-6 | ORF6-7 |  |  |  |  |  |  |  |  |  |
| Pool-O-5 | ORF7a-1 | ORF7a-2 | ORF7a-3 | ORF7a-4 | ORF7a-5 | ORF7a-6 | ORF7a-7 | ORF7a-8 | ORF7a-9 | ORF7a-10 | ORF7a-11 | ORF7a-12 | ORF7a-13 | ORF7a-14 | ORF7a-15 |  |
| Pool-O-6 | ORF8-1 | ORF8-2 | ORF8-3 | ORF8-4 | ORF8-5 | ORF8-6 | ORF8-7 | ORF8-8 | ORF8-9 | ORF8-10 | ORF8-11 | ORF8-12 | ORF8-13 | ORF8-14 | ORF8-15 | ORF8-16 |
| Pool-O-7 | Env-1 | Env-2 | Env-3 | Env-4 | Env-5 | Env-6 | Env-7 | Env-8 | Env-9 |  |  |  |  |  |  |  |
| Pool-O-8 | M-1 | M-2 | M-3 | M-4 | M-5 | M-6 | M-7 | M-8 | M-9 | M-10 | M-11 | M-12 | M-13 | M-14 | M-15 | M-16 |
| Pool-O-9 | M-17 | M-18 | M-19 | M-20 | M-21 | M-22 | M-23 | M-24 | M-25 | M-26 | M-27 | M-28 | M-29 |  |  |  |
| Pool-O-10 | NP-1 | NP-2 | NP-3 | NP-4 | NP-5 | NP-6 | NP-7 | NP-8 | NP-9 | NP-10 | NP-11 | NP-12 | NP-13 | NP-14 | NP-15 | NP-16 |
| Pool-O-11 | NP-17 | NP-18 | NP-19 | NP-20 | NP-21 | NP-22 | NP-23 | NP-24 | NP-25 | NP-26 | NP-27 | NP-28 | NP-29 | NP-30 | NP-31 | NP-32 |
| Pool-O-12 | NP-33 | NP-34 | NP-35 | NP-36 | NP-37 | NP-38 | NP-39 | NP-40 | NP-41 | NP-42 | NP-43 | NP-44 | NP-45 | NP-46 | NP-47 | NP-48 |
| Pool-O-13 | NP-49 | NP-50 | NP-51 | NP-52 | NP-53 | NP-54 | NP-55 | NP-56 | NP-57 | NP-58 | NP-59 |  |  |  |  |  |

**Supplementary Table 2:**
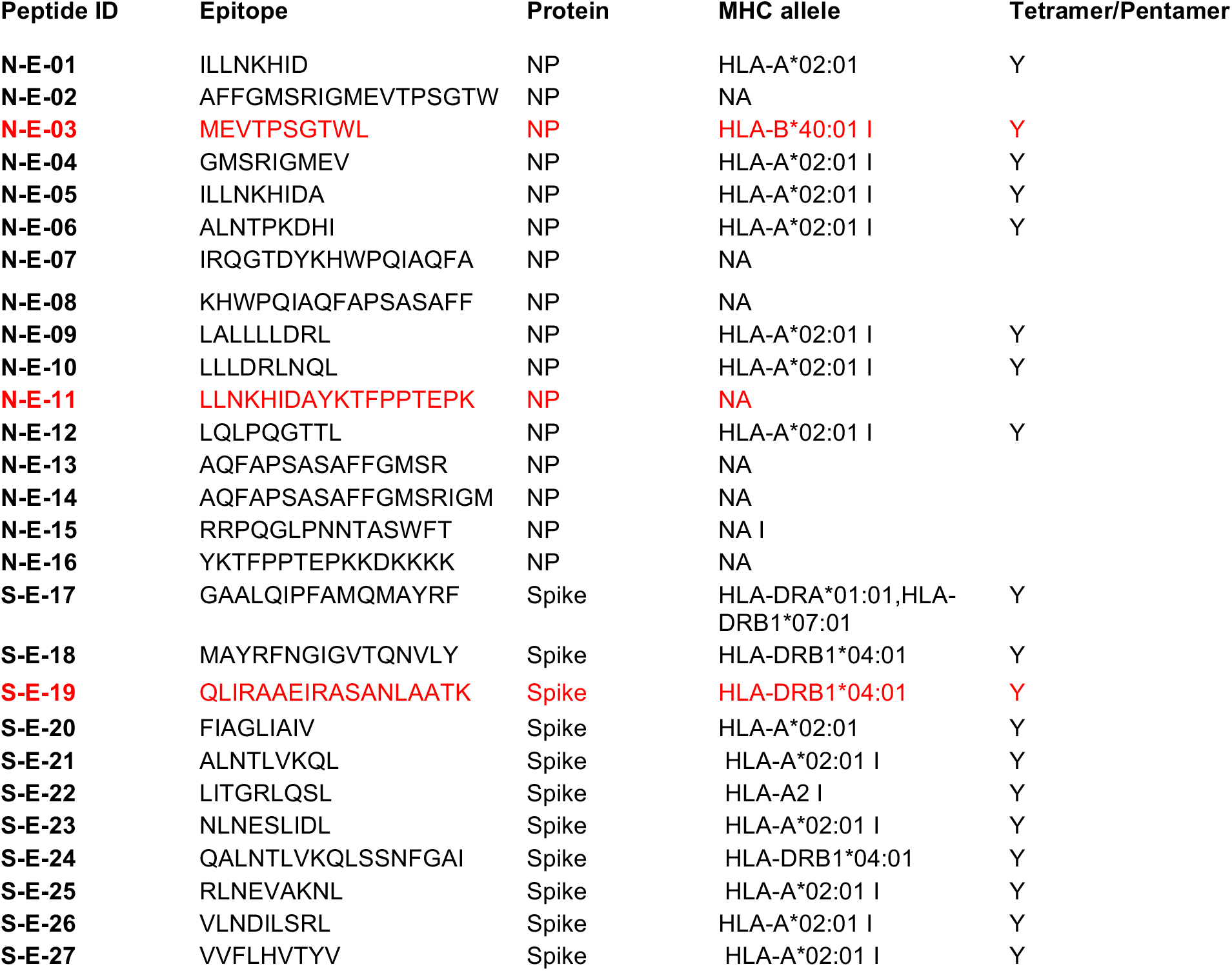
Known SARS epitopes with identical sequences to SARS-CoV-2, and Tetramers/Pentamers. Red highlights the epitope responses detected in the patients recovered from COVID-19.

**Supplementary Table 3:**
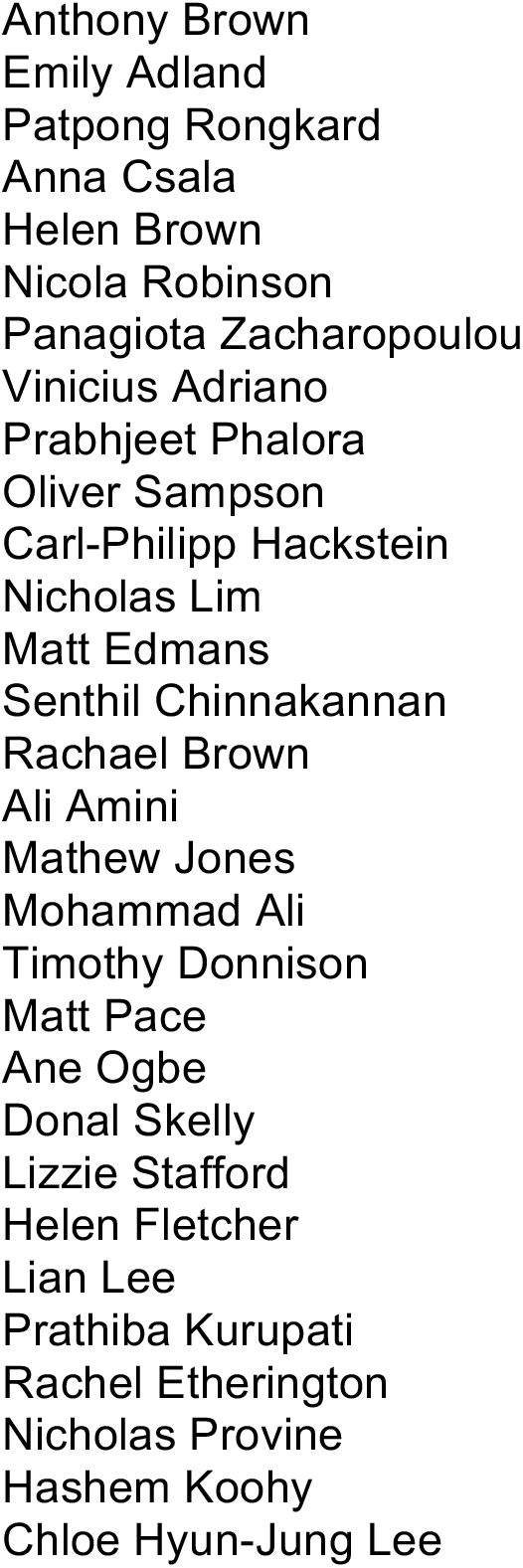
Oxford Immunology Network Covid-19 response: T cell Immunity Team – additional contributors

